# Evolution towards extremely high β-lactam resistance in *Mycobacterium abscessus* outbreak strains

**DOI:** 10.1101/2024.05.08.593223

**Authors:** Eva le Run, Hervé Tettelin, Steven M. Holland, Adrian M. Zelazny

## Abstract

Treatment of *Mycobacterium abscessus* pulmonary disease requires multiple antibiotics including intravenous β-lactams (e.g., imipenem, meropenem). *M. abscessus* produces a β-lactamase (Bla_Mab_) that inactivates β-lactam drugs but less efficiently carbapenems. Due to intrinsic and acquired resistance in *M. abscessus* and poor clinical outcomes, it is critical to understand the development of antibiotic resistance both within the host and in the setting of outbreaks.

We compared serial longitudinally collected *M. abscessus* subsp. *massiliense* isolates from the index case of a CF center outbreak and four outbreak-related strains. We found strikingly high imipenem resistance in the later patient isolates, including the outbreak strain (MIC >512 µg/ml). The phenomenon was recapitulated upon exposure of intracellular bacteria to imipenem. Addition of the β-lactamase inhibitor avibactam abrogated the resistant phenotype. Imipenem resistance was caused by an increase in β-lactamase activity and increased *bla*_Mab_ mRNA level. Concurrent increase in transcription of preceding *ppiA* gene indicated upregulation of the entire operon in the resistant strains.

Deletion of the porin *mspA* coincided with the first increase in MIC (from 8 to 32 µg/ml). A frameshift mutation in *msp2* responsible for the rough colony morphology, and a SNP in ATP-dependent helicase *hrpA* co-occurred with the second increase in MIC (from 32 to 256 µg/ml). Increased Bla_Mab_ expression and enzymatic activity may have been due to altered regulation of the *ppiA*-*bla*_Mab_ operon by the mutated HrpA alone, or in combination with other genes described above. This work supports using carbapenem/β-lactamase inhibitor combinations for treating *M. abscessus*, particularly imipenem resistant strains.

## Introduction

*Mycobacterium abscessus* is a pathogenic, multidrug-resistant organism within the rapidly growing nontuberculous mycobacteria (NTM) that causes lung, soft tissue, and disseminated infections. Risk factors include preexisting lung conditions and immunodeficiency. Recent studies have shown that the incidence and prevalence of NTM infections are on the rise in the United States (1) (2). Lung disease accounts for the majority of NTM infections (3) in susceptible individuals such as those with cystic fibrosis (CF). According to the CF Foundation, 10% of CF patients presented with positive cultures for *M. abscessus* (4).

Notorious outbreaks of *M. abscessus* have occurred in CF Centers worldwide including Seattle, WA, USA (5) and Papworth, UK (6). Genomic analysis by Tettelin *et al*. showed high similarity among strains from the four patients involved in the Seattle outbreak but also a surprisingly high-level relatedness with isolates from geographically distant outbreaks (7). Genomic comparison of strains collected serially from CF patients and from different CF patients over time have allowed inferences on patient-to-patient transmission (6).

Recommended treatment for pulmonary infection by *M. abscessus* consists of an initial phase of IV and oral drugs including a macrolide, an aminoglycoside, a β-lactam (imipenem or cefoxitin) and a glycylcycline for 3 months followed by continuation phase of four to five oral drugs for at least 14 months (1,8). Unfortunately, *M. abscessus* is naturally resistant to many antibiotics by different mechanisms (9). Most strains display resistance to macrolides through the inducible *erm41* gene (for *M. abscessus* subsp. *abscessus* and *bolletii*). In addition, strains may acquire macrolide resistance through specific mutations in the 23S rRNA gene.

The β-lactam imipenem is often used intravenously during the initial phase of treatment of *M. abscessus*. Several studies discussed the use of imipenem in combination with other drugs including β-lactamase inhibitors (10,11), other β-lactam drugs (12,13) and other antibiotics (14,15) for the treatment of *M. abscessus* infections. Imipenem was strongly associated with treatment success for *M. abscessus* pulmonary disease in a meta-analysis study (adjusted odds ratio 2.65, 95% CI 1.36–5.10) (16). As a β-lactam, imipenem covalently binds to penicillin binding proteins (PBPs) responsible for the synthesis of peptidoglycan. *Mycobacterium abscessus* has two major L,D-transpeptidases responsible for the synthesis of the peptidoglycan (17); only one of them is inhibited by carbapenems (18).

Resistance to β-lactams is mediated by the production of a class A β-lactamase encoded by the *bla*_Mab_ gene (MAB_2875). Bla_Mab_ can efficiently hydrolyze penams (amoxicillin, ampicillin) and cephems *in vitro*. However, imipenem and cefoxitin, both part of the recommended treatment for *M. abscessus*, are hydrolyzed at a lower rate than other β-lactam drugs (19). β-lactamase inhibitors such as avibactam have shown promising results for *M. abscessus* when used in combination with imipenem (20,21). Avibactam improved the efficacy of imipenem against *M. abscessus in vitro* as well as in macrophages and zebrafish embryos (22).

As part of the investigation of the Seattle outbreak, we obtained serial isolates of *M. abscessus* subsp *massiliense* (hereafter referred to as *M. massiliense*) from the index case over eight years, including the outbreak strain. Recently, we described the development of clarithromycin and amikacin resistance in those organisms (23). Notwithstanding, we speculated that this organism could have developed resistance to additional antimicrobials, as part of its evolution within the patient, culminating in the outbreak strain.

Using phenotypic and genomic analysis of serial isolates from the Seattle outbreak index patient and secondary cases, we describe the development of extremely high-level imipenem resistance and provide insights into the resistance mechanism.

## Results

### Imipenem activity against patient 2B serial isolates

Serially collected isolates from patient 2B were screened for differences in susceptibility to imipenem. MICs of 11 isolates and type strain *M. massiliense* CCUG48898^T^ were determined by broth microdilution method in 7H9sB media (Table 1). The MIC of imipenem against isolate 2B1 to 2B4 was 8 µg/ml, comparable to that of CCUG48898^T^ (16 µg/ml). There was a slight increase in MIC in strains 2B5 and 2B6 (32 µg/ml). Remarkably, isolates 2B7 to 2B11 showed a dramatic increase in MIC beyond the limit of quantification of the assay (> 512 µg/ml). This rise in MIC coincided with the shift from smooth to rough colony morphotype (first seen with strain 2B7). Based on these results, we selected three representative strains 2B1, 2B5 and 2B11 to further investigate the mechanism of resistance.

**Table 1.**
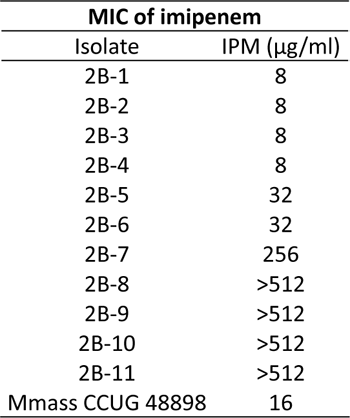
Minimal Inhibitory Concentrations of imipenem against 2B patient isolates and *M. massiliense* CCUG 48898T. MICs were determined using the broth microdilution method in 7H9sB. MICs were read after 48 hrs incubation at 37°C and are expressed in µg/ml. Values are the median of five independent replicates.

| MIC of imipenem |  |
| --- | --- |
| Isolate | IPM (µg/ml) |
| 2B-1 | 8 |
| 2B-2 | 8 |
| 2B-3 | 8 |
| 2B-4 | 8 |
| 2B-5 | 32 |
| 2B-6 | 32 |
| 2B-7 | 256 |
| 2B-8 | >512 |
| 2B-9 | >512 |
| 2B-10 | >512 |
| 2B-11 | >512 |
| Mmass CCUG 48898 | 16 |

### Imipenem activity against intracellular *M. massiliense* in THP1 macrophages

To evaluate antimicrobial resistance of our strains within the intracellular environment, THP1 cells differentiated into macrophages were infected with *M. massiliense* CCUG48898^T^, 2B1 or 2B11 strains and treated with imipenem for 48 hours (24). Fold changes, expressed as colony forming unit (cfu) counts after 48 hours of treatment over initial cfu count, are shown for each condition in Figure 2. Fold change values above 1 represent growth of intracellular bacteria, while those below 1 indicate bacterial killing. All three strains showed similar growth inside macrophages in the absence of imipenem (Mass at 50.5, 2B1 at 52.2 and 2B11 42.6, respectively). When treated with imipenem at 32 µg/ml, the fold change of CCUG48898^T^ was 0.4, demonstrating intracellular killing. Strain 2B1 partially withstood imipenem treatment showing a 1.5-fold change, even though the difference was not statistically significant.

**Figure 1.**
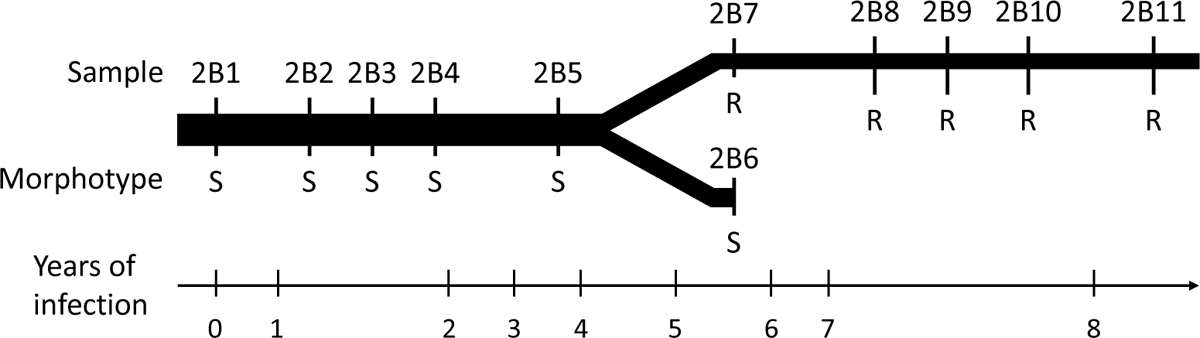
Timeline of serial isolates from patient 2B. Patient 2B had cystic fibrosis complicated by a long-term respiratory infection with *M. abscessus* subsp. *massiliense*. Isolates were obtained over an 8-year period until patient’s death. s: smooth colony morphology, R rough colony morphology.

**Figure 2.**
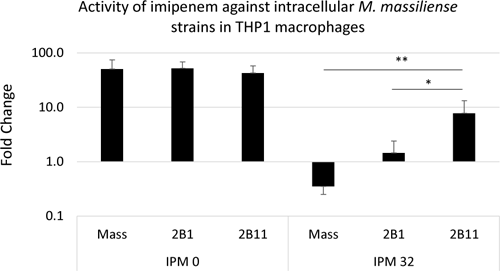
Activity of imipenem against intracellular *M. massiliense* strains in THP1 macrophages. PMA differentiated THP1 cells were infected by different *M. massiliense* strains, then treated with <u>or without</u> imipenem at 32 ug/ml for 24 hrs or left untreated. Cells were then lysed and intracellular bacterial load was determined. Fold change are calculated by divided the cfu count at 48 hrs by the initial bacterial load before treatment. Fold change >1 corresponds to intracellular growth of the bacteria, Fold change <1 reflects the killing of intracellular bacteria.. *P<0.05, **P<0.01, T-test analysis

Remarkably, strain 2B11 showed a 7.8-fold change with imipenem, highlighting the remarkably poor activity of the drug against intracellular 2B11. These data confirmed the higher level of resistance of 2B11 to imipenem compared to the initial infectious strain 2B1 and CCUG48898^T^ in the context of infected macrophages.

### β-lactamase enzymatic activity assay

*Mycobacterium massiliense* produces a class A β-lactamase, Bla_Mab_, responsible for the hydrolysis of β-lactams, including imipenem. In order to investigate if the observed differences in MIC could be due to changes in β-lactamase activity, we measured the specific enzymatic activity in protein extracts from each strain with the chromogenic substrate nitrocefin. As seen in Figure 3, strain 2B1 showed baseline β-lactamase activity (26.3 nmol/min/mg) which was comparable to that of *M. massiliense* CCUG48898^T^ control (35 nmol/min/mg). Interestingly, 2B5 β-lactamase activity was 2.5 times higher than that of 2B1 (64 nmol/min/mg, *p*<0.05), while the enzymatic activity of 2B11 increased even further, to 4 times that of 2B1 (108.9 nmol/min/mg, *p*<0.05). These results provide an explanation for the rise in imipenem resistance in the late isolates of the patient.

**Figure 3.**
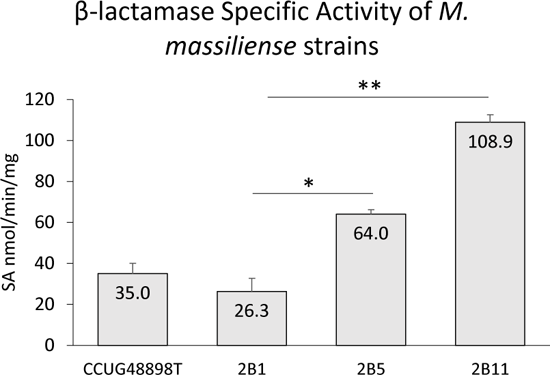
β-lactamase specific activity of 2B patient isolates and *M. massiliense* CCUG48898T. specific activity (sA) was determined by following the hydrolysis of nitrocefin by spectrophotometry (λ = 486 nm). Data shown are the mean and standard deviation of five independent replicates. *P<0.05, **P<0.01, T-test analysis.

### Transcription level of *bla*_Mab_ gene assay

We sought to determine whether the higher β-lactamase activity observed in isolate 2B11 was due to enhanced enzymatic activity or increased expression of Bla_Mab_. We quantified *bla*_Mab_ mRNA and *dna*K housekeeping gene control by RT-PCR calculated versus those of CCUG48898^T^ control strain (RQ).

Since *bla*_Mab_ is the second gene in an operon (Figure 4A) we also measured the mRNA level of the preceding gene, *ppiA*. Results for *ppiA* were reported the same way as for *bla*_Mab_. As shown in Figure 4B, 2B1 and 2B5 strains showed baseline levels of *bla*_Mab_ mRNA which were similar to that of CCUG48898^T^ control (1.2 and 1.3 respectively). Interestingly, the *bla*_Mab_ transcription level of strain 2B11 was 2.6 times higher than that of 2B1, 2B5 and CCUG48898^T^. Thus, the increase in *bla_Mab_ mRNA* levels in strain 2B11 correlated with the higher Bla_Mab_ activity described above. The transcription level of *ppiA* was significantly increased in 2B11 compared to 2B1 and 2B5 (10.84 vs. 1.80 and 2.72, respectively; Figure 4C). Concurrent increases in *ppiA* and *bla*_Mab_ transcription indicate upregulation of the entire operon in the resistant strains.

**Figure 4.**
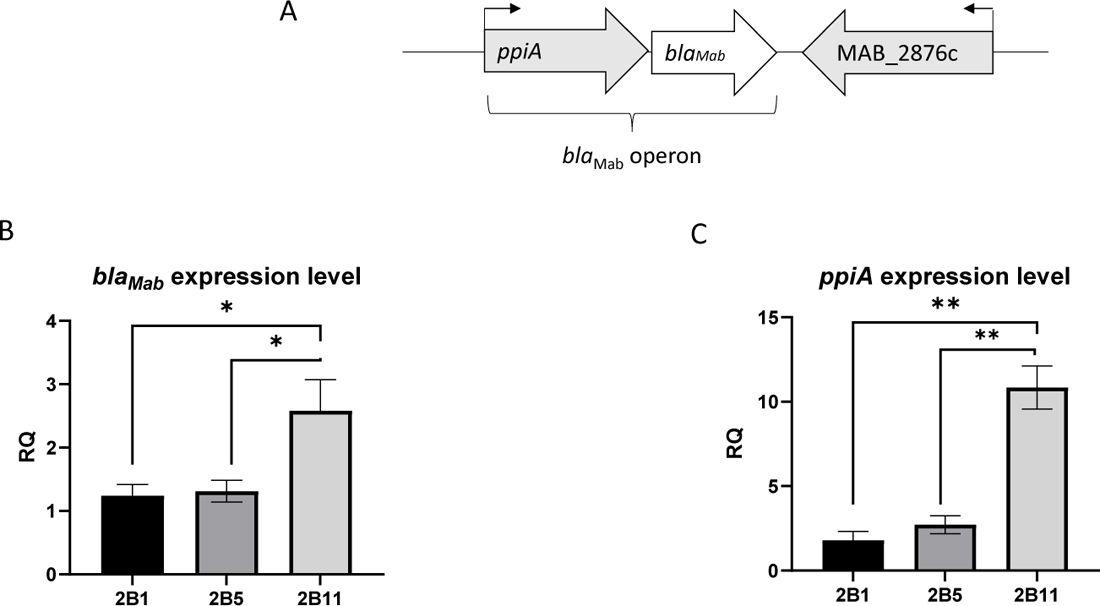
Expression level of *ppiA-bla*_Mab_ operon in patient 2B i (A) schematic representation of *ppiA*-*bla*_Mab_ operon. The mRNA level of *bla*_Mab_ (B) and *ppiA* (C) were determined by RT-Q-PCR using *dnaK* as housekeeping gene and calculated versus those of CCUG48898T control strain (RQ). Data shown are the mean and standard deviation of three independent replicates. *P<0.05; **P<0.01 using T-test.

### Inhibition of Bla_Mab_ activity by avibactam

Avibactam is a second generation β-lactamase inhibitor from the diazabicyclooctanes (DBO) family that was shown to inhibit Bla_Mab_ *in vitro* (20). We determined MIC of imipenem in combination with 4 µg/ml of avibactam. As shown in Table 2, addition of avibactam led to a significant decrease in MIC against all tested strains. MIC against 2B1 and CCUG48898^T^ decreased two and four-fold, respectively, to 4 µg/ml. The MIC against 2B5 decreased by 2-fold. Notably, avibactam led to a striking reduction in MIC of strain 2B11 (from > 512 to 32 µg/ml).

**Table 2.** Minimal Inhibitory Concentrations of imipenem alone or in combination with avibactam against 2B patient isolates and *M. massiliense* CCUG48898T. MICs were determined using the broth microdilution method in 7H9sB. Different concentrations of imipenem were tested in combination with 4 µg/ml of avibactam. MICs were read after 48 hrs incubation at 37°C and are expressed in µg/ml. Values are the median of five independent replicates.

| Isolate | IPM (µg/ml) | IPM + Avibactam (µg/ml) |
| --- | --- | --- |
| 2B-1 | 8 | 4 |
| 2B-5 | 32 | 8 |
| 2B-11 | >512 | 32 |
| Mmass CCUG48898 | 16 | 4 |

### Imipenem activity against the Seattle CF outbreak strains

Strain 2B11 was identified to be part of a global dominant circulating clone of *M. massiliense* responsible for outbreaks in CF centers. We obtained four strains from other patients in the Seattle CF outbreak. Imipenem MICs were performed against strains G_2446, G_2258 G_2455 and G_2272, all of which were previously found to be closely related by genomic sequencing. As shown in (Table 3), G_2272 and G_2446 showed imipenem MICs of 512 µg/ml and above, respectively. Imipenem MIC against G_2455 was 128 µg/ml. Surprisingly, MIC against G_2258 was only 8 µg/ml, comparable to that of the early isolates from the 2B patient. We performed further genomic comparisons between G_2258 and 2B11 and the other outbreak strains to identify genetic changes that could explain the variations in imipenem resistance.

**Table 3.** Minimal Inhibitory Concentrations of imipenem against outbreak isolates collected in seattle, UsA. MICs were determined using the broth microdilution method in 7H9sB. MICs were read after 48 hrs incubation at 37°C and are expressed in µg/ml. Values are the median of five independent replicates.

| Imipenem MICs |  |
| --- | --- |
| Isolate | IPM (µg/ml) |
| G_2258 | 8 |
| G_2272 | 512 |
| G_2446 | >512 |
| G_2455 | 128 |

### Mutations concurring with the development of moderate and extremely high imipenem resistance

Following whole genome sequencing using Illumina platform reported previously, we performed long-read sequencing of strains 2B1, 2B5, 2B7 and 2B11 (23). The 4 complete, gap-free genome sequences were compared with genome sequences of other strains from the same outbreak (Table S3 for accession numbers) using Sybil (25,26) and genomic regions spanning clusters of orthologous genes carrying changes between isolates were realigned using Muscle (27).

This analysis revealed several mutations that appeared concurrently with the development of imipenem resistance, shown in Table 4. A mutation in *mspA* (porin) coincided with the first increase in MIC in 2B5 (from 8 to 32 µg/ml). The second increase in MIC (from 32 to 256 µg/ml) was detected in 2B7 which was concurrent with a frameshift mutation in *msp2,* disrupting the glycopeptidolipid (GPL) biosynthesis pathway and changing the colony morphology from smooth to rough, and a SNP in the ATP-dependent helicase *hrpA*. All three mutations persisted in 2B11 (MIC >512 µg/ml), which caused the Seattle CF outbreak.

**Table 4.** Mutations co-occurring with acquisition or loss of the resistance phenotype. Mutations were identified by a using data from long read sequencing data (this manuscript) combined with illumina data (previously published). The appearance of the first three mutations coincide with resistance phenotype development. The four mutation coincides with loss of resistance in one of the outbreak strains.

| Gene Name | Description | GO Biological process/Function | MAB Locus | Amino acids (in isolate) | In 2B isolates | Outbreak strains | Process | Potential explanations |
| --- | --- | --- | --- | --- | --- | --- | --- | --- |
| *mspA* | porin | Porin activity | MAB_1080 | 224 (2B-1)<br>82 (2B-11) | ORF truncated from isolates 2B-5 to 2B-11 | Identical to 2B11 | Truncation<br>Deletion of a piece of sequence | appears with the first increase in MIC (from 8 to 32 µg/ml) |
| *mps2* | Non-ribosomal peptide synthetase | Glycopeptidolipid (GPL) production | MAB_4098c | 2582 (2B-1)<br>267 and 2327 (2B-11) | 2 ORFs in isolate 2B-7, 2B-9, and 2B-11 | Identical to 2B11 | Frameshift<br>Deletion of one A<br>(G_2258 also has an additional deletion of a C further down, G_2246 also has an additional deletion of a G further down) | appears with the second increase in MIC (from 32 to 256 µg/ml).<br>Leads to the R morphotype. |
| *hrpA* | ATP-dependent RNA helicase | nucleic acid metabolic process | MAB_0056c | 1286 | G->T in 2B7, 2B9, 2B11. | Identical to 2B11 | N446K | appears with the second increase in MIC (from 32 to 256 µg/ml) |
| *ppiA* | probable peptidyl prolyl cis trans isomerase | Cis/trans isomerization of prolyl bonds | MAB_2874 | 307aa (2B1-2B11)<br>597aa (G_2258) | - | 2 ORFS fused in G_2258 | Frameshift<br>Deletion of one C | appears with the loss of imipenem resistance in outbreak strain G_2258.<br>leads to the fusion of *ppiA* and *BlaMab* (MAB_2875) linked by random amino acids |

We also compared the genomes of the four additional strains from the Seattle outbreak. All four strains shared the same frameshift mutation in *msp2* and SNP in *hrpA* genes. However, G_2258 and G_2246 presented a second mutation downstream in *msp2* (deletion of C and G, respectively), that did not restore the frameshift of the first mutation.

Interestingly, outbreak strain G_2258, which displayed a low imipenem MIC (8 µg/ml) despite being related to the highly resistant strain 2B11, showed a single nucleotide deletion in *ppiA* (MAB_2874). In the *M. abscessus* genome, this gene precedes MAB_2875 encoding the β-lactamase, Bla_Mab,_ and is part of the same operon. This deletion caused a frameshift resulting in a fusion protein from both genes, albeit with a missense amino acid sequence. This can explain the loss of imipenem resistance in outbreak strain G_2258 Table 3.

## Discussion

Using serial isolates from a CF patient over eight years, we were able to track and characterize the development of extraordinarily high-level imipenem resistance in *M. massiliense* from the first isolate 2B1 (MIC = 8 µg/ml), to isolate 2B5 (MIC= 32 µg/ml) through the last isolate 2B11 (MIC >512 µg/ml). The patient was the index case of a well-documented CF center outbreak in Seattle, WA (5).

A review of *M. abscessus* susceptibility data from our laboratory in the past two years found that 62% and 32% of tested strains were intermediate (MIC of 8-16 µg/ml) and resistant (MIC ≥32 µg/ml) to imipenem, respectively. Multiple publications described the MIC90 of imipenem being at or above 64 µg/ml in clinical strains (Fröberg *et al.* used 1014 samples (28), Ying *et al*. used 46 samples (29) and Liu*, et al*. used 114 samples (30)). Of note, most clinical microbiology laboratories use commercial antimycobacterial susceptibility testing plates for MIC determinations. The highest imipenem concentration tested in the commonly used RAPMYCO and RAPMYCO2 Sensititre panels is 64 and 32 µg/ml, respectively, which takes into consideration CLSI’s breakpoint of ≥32 µg/ml for imipenem resistance. Since this commercial panel only tests a limited range of MICs, there is scant information on the occurrence and frequency of high-level imipenem resistance in *M. abscessus*.

Using a host cell-mycobacteria infection model with the macrophage-differentiated THP1 cell line (24,31), we showed that the high level resistance phenotype was recapitulated upon exposure of intracellular bacteria to imipenem. While 32 µg/ml of imipenem showed killing activity against *M.*. *massiliense* type strain CCUG48898^T^ after 48 hrs, this effect was reduced for the first clinical strain 2B1, although not significantly (0.4 ±0.1 vs 1.5 ± 0.9; figure 2). Remarkably, the last clinical strain 2B11 continued replicating within macrophages despite the antibiotic treatment (fold change of 7.8 ± 5.5) underscoring its high level of imipenem resistance.

*Mycobacterium abscessus* Bla_Mab_ displays a low catalytic efficiency against imipenem. In Dubée *et al,* imipenem showed an MIC of 8 µg/ml against *M. abscessus* type strain ATCC 19977, which is in the susceptible category according to CLSI guidelines. The MIC decreases to 4 µg/ml upon deletion of the *bla*_Mab_ gene (strain *M. abscessus* Δ*bla*_Mab_) (20). We used a nitrocefin hydrolysis kinetics assay to assess specific β-lactamase enzymatic activity in our *M. abscessus* subsp *massiliense* strains. The first strain 2B1 showed similar β-lactamase activity to that of the type strain control CCUG48898^T^. Notably, the β-lactamase activity of the last strain 2B11 was about 4 times higher (Figure 3). This increased Bla_Mab_ activity could explain the high-level imipenem resistance observed in 2B11.

To further characterize the resistance phenotype, levels of *bla*_Mab_ mRNA transcripts were measured by RT Q PCR. Strain 2B11 showed 2.2 times higher levels of *bla*_Mab_ mRNA compared to 2B1 (Figure 4), which correlated with the increased β-lactamase activity and the extremely high MIC observed in the resistance strain. In addition, it showed an increased expression of *ppiA*, a gene immediately upstream of *bla*_Mab_, and under the same promoter. Similar results were obtained for the first highly resistant strain, 2B7 (data not shown). We speculated whether the increase in β-lactamase mRNA and enzymatic activity alone could account for the extremely high imipenem MIC or was it multifactorial. To address this question, we assessed imipenem activity against the strains with and without the β-lactamase inhibitor avibactam.

Avibactam, a 2^nd^ generation diazabicyclooctane (DBO) β-lactamase inhibitor, has been previously shown to inhibit Bla_Mab_ (32) and rescue β-lactam activity *in vitro*, in intracellular infection models, and in animal models (20,33). Commercially, imipenem is available in combination with another DBO, relebactam. Imipenem/relebactam and imipenem/avibactam showed similar activity against *M. abscessus* ATCC19977 *in vitro* and in macrophage model (31,34). Avibactam reduced imipenem MIC against *M. abscessus* ATCC19977 to comparable levels of the *bla*_Mab_ gene deleted strain (*M. abscessus* Δ*bla*_Mab_) and restored the ability of imipenem to kill intracellular mycobacteria. Similarly, in our study, the addition of avibactam significantly decreased the imipenem MIC of our highly resistant strains to levels closer to those of the early susceptible ones (Table 2). These results support the hypothesis that the higher transcription level of the *bla*_Mab_ gene and subsequent increased enzymatic activity of Bla_Mab_ is the main (if not the only) driver of the extraordinarily high imipenem MICs observed in our clinical strains. Our work further highlights the value of β-lactam/β-lactamase inhibitor combinations in treatment regimens for *M. abscessus* infections.

Genomic comparisons between the first isolate (2B1), the moderately resistant isolate (2B5), the first extremely high resistant isolate (2B7) and the last isolate (2B11) were carried out to look for mutations or rearrangements that might explain these findings. We focused on those gene changes which appeared in 2B5 and 2B7 and persisted throughout the later isolates. Mutations occurred in genes involved in cell envelope, transcriptional regulation, and lipid and amino acid metabolism (Table 4).

Sequencing of 2B isolates proved a large deletion of the *mspA* gene (MAB_1080) from 2B5 through 2B11, coinciding with the first increase in imipenem resistance. It has been shown in *M. smegmatis* that MspA is the major porin protein (35). Previously, we described the effects of this deletion, which included a slower growth rate, explained by lower nutrients (eg., glucose) intake. *M. abscessus* with a mutated *mspA* showed an increase in virulence and inflammation (23,36). Deletion of porin genes from *M. massiliense* CCUG48898^T^ had no effect on the antibiotic susceptibility profile (36). In contrast, Danilchanka *et al*, reported higher quinolone and β-lactams resistance in *M. smegmatis* strains lacking *mspA* (37). Thus, it remains unclear whether changes in the *mspA* gene were responsible for the initial increase in imipenem resistance observed in strains 2B5 and 2B6.

Loss of GPL production through mutations in GPL pathway genes including *mps2* (MAB_4098c) is responsible for the rough (R) colony morphology in *M. abscessus* (38–40). We showed that the dramatic increase in imipenem MIC observed in strain 2B-7 through 2B11 coincided with the switch from smooth to rough morphotype. Different studies have tried to establish a link between colony morphotype and resistance to common antibiotics. Lavollay *et al*. (41) linked the R morphotype of clinical strains to higher imipenem and cefoxitin MICs. In contrast, Hershko *et al*. (42) used transposon technology to generate a rough GPL-defective strain and showed that the loss of MAB_4099c led to only a minor (one dilution) increase in the MIC of imipenem.

A single point mutation was found in *hrpA* (MAB_0056c) in 2B7 and conserved in all later isolates. *hrpA* encodes for an ATP RNA helicase containing a DEAH box domain (43). This protein has not been extensively studied in mycobacteria. However, in *Neisseria meningitidis* (44), it is involved in biofilm formation, while in *E. coli* it is necessary for mRNA processing of the fimbrial operon (45). HrpA is also important in gene regulation of *Borrelia burgdorferi* at a post-transcriptional level and essential for virulence in mice (46,47).

Using the database STRING, ten proteins have been identified as interacting with HrpA, one of them being PpiA, a peptidyl polyl cis trans isomerase which precedes the β-lactamase gene *bla*_Mab_ in an operon (data not shown). We showed in this study that transcription and expression of both *ppiA* and *bla*_Mab_ were increased in the highly resistant strains 2B7 and 2B11 (Figure 4). We hypothesize that *HrpA* is involved in the regulation of *ppiA* at the RNA level, which subsequently affects Bla_Mab_ expression as part of the same operon. The *hrp*A mutation and subsequent N446K amino acid change observed in strains 2B7 and 2B11 could alter HrpA regulation of the *ppiA*-*bla*_Mab_ operon leading to a higher expression of Bla_Mab_ responsible for the extremely high resistance to imipenem.

Occurrence of extremely high-level resistance to imipenem is alarming, particularly in this case as the strains were clinical isolates from a CF patient that led to a serious outbreak. Transmission of a strain resistant to multiple antibiotics to patients who have not been treated with those drugs is a major concern (6).

Using serial isolates from a CF patient, we followed and dissected the development of extremely high-level resistance to imipenem appearing progressively over time in strains of *M. massiliense* that were involved in a CF outbreak. A significant increase in transcription level and enzymatic activity of β-lactamase observed in the resistant strains accounted for the resistance phenotype. While no mutations were found on the β-lactamase *bla*_Mab_ gene or its promoter, we hypothesize that increased Bla_Mab_ expression and enzymatic activity could be due to the observed mutation in *hrp*A alone or in combination with mutations in the additional genes described above. Our findings also re-affirm the importance of the addition of a β-lactamase inhibitor to the common antibiotic treatment of *M. abscessus* to prevent the β-lactamase from degrading imipenem. The imipenem/relebactam combination is available clinically and should be considered for the treatment of *M abscessus*, particularly for imipenem resistant strains.

## Material and Methods

### Strains & media

Serial isolates of *M. massiliense* from the CF patient (referred to as patient 2B), and the Seattle outbreak strains were stored at – 80°C. Information on the strains including name, date of isolation and GenBank accession numbers is provided in Supplementary table 1 and a previous publication (23). *M. abscessus* subsp. *massiliense* CCUG48898^T^ was used as reference strain.

Strains were subcultured from frozen stocks onto 7H11 agar plates and incubated for four days at 37°C, 5% CO_2_. Liquid media cultures were performed from single colonies in 7H9 supplemented with ADC and 1% glycerol (7H9sB) for three days at 37°C with shaking. Cultures were diluted to OD600 0.05 in 7H9sB at 37°C and grown until exponential phase (OD600 ∼0.7-1.0) prior to experiments.

Human leukemia monocytic cell line THP1 cells were grown from frozen stocks in RPMI supplemented with HEPES 10mM, Sodium pyruvate 1mM and 10% Fetus Bovine Serum (RPMIsB) and incubated at 37°C 5% CO_2_. All cell experiments were performed using RPMIsB medium.

### MICs testing

Minimal inhibitory concentrations (MICs) were determined by the broth microdilution method. Bacteria grown in 7HPsB for 4 days were diluted to an OD600 = 0.05 and incubated for 24 hrs at 37°C with shaking at 180 rpm. These cultures were used to inoculate 96-well round-bottom plates (∼5.10^5^ cfu/ml) containing 2-fold dilutions of antibiotics in 7H9sB. Plates were examined for bacterial growth after 48 hrs incubation at 37°C. The MIC was defined as the lowest drug concentration inhibiting visible growth. Experiments were performed in quintuplets and data shown is the median in µg/ml. Imipenem was purchased from Sigma-Aldrich. Antibiotic solutions were made fresh prior to each replicate.

### Intracellular killing of *M. massiliense* in differentiated THP1 macrophages

THP-1 cells were differentiated into 12-well plates (5 × 10^5^ cells per 1 mL well), with PMA 20 ng/ml in RPMIsB for 24 hrs at 37°C 5% CO_2_. Cells were then infected with *M. massiliense* CCUG48898^T^, 2B1 or 2B11 strains at MOI 10:1 for 3 hrs. at 37°C, 5% CO_2_. Imipenem (32 µg/ml) was then added to the treatment wells and plates incubated for 48 hrs. Medium with or without imipenem was renewed every 24 hrs. Quantitative cultures of intracellular bacteria were performed by plating serial dilutions of macrophage lysates on LB agar plates incubated for 4 days at 37°C. Results represent the means (±SD) of five independent experiments.

### β-lactamase enzymatic activity

β-lactamase specific activity of total protein extract of *M. massiliense* cultures was measured using the chromogenic β-lactam substrate nitrocefin. Briefly, strains were grown in 7H9sB until OD600 ∼1. Five ml of cultures were spun down and the bacterial pellet resuspended in 1 ml PIPES 2mM pH 6.8 containing zirconia/silica beads 0.1 mm diameter (BioSpec Product). Bacteria were then mechanically disrupted using a bead beater homogenizer (PowerLyzer 24, Qiagen; 2 cycles of 45s 5000 rpm / 5 min ice). After an additional 5 min on ice, samples were spun down and supernatants containing total soluble proteins removed and stored at – 20°C until use. Prior to experiments, samples were thawed and 10 µl of total protein extract was added to 100 µM nitrocefin (Sigma-Aldrich) solution in PIPES 2 mM pH 6.8 and substrate hydrolysis measured spectrophotometrically over 30 minutes (Cytation5, Biotek) at 486 nm at 20°C. In addition, protein concentration was quantified by the BCA assay (Pierce^TM^ BCA Protein Assay kit, Thermofisher). Specific activity was calculated as the concentration of hydrolyzed nitrocefin produced per minute per mg of total protein extract. Results represent the mean values expressed in nmol/min/mg of five independent experiments.

### RNA extraction and quantification

RNA was extracted from each strain cultivated in 7H9sB at 37°C for two days using Nucleospin kit (Macherey-Nagel, Allentown, PA) following manufacturer instructions with few modifications. Five ml of the same cultures used for protein extraction were spun down and resuspended in 350 µl of RA1 lysis buffer. Bacteria were resuspended with zirconia/silica beads 0.1 mm diameter and disrupted mechanically (PowerLyzer 24 Qiagen) two cycles of 45s 5000 rpm with 5 minutes ice in between. After 5 minutes on ice, tubes were spun down and the supernatant was transferred to another tube and 3.5 µl of β-mercaptoethanol were added. RNA was obtained in RNase/DNase free water and the concentration measured with Nanodrop (Thermofisher Scientific). RT-Q-PCR was performed with the Power SYBR™ Green RNA-to-CT™ 1-Step Kit (Thermofisher Scientific). Primers were designed to target different genes of interests (e.g. *ppiA*, *bla_Mab_, dnaK*) (Supplementary table 2).

### Long Read sequencing

For genomic DNA (gDNA) extraction, strains 2B1, 2B5, 2B7 and 2B11 were cultured on 7H11 plates at 37°C for 4 days. Bacterial colonies were resuspended in 0.05 M Tris HCl – 10 mM EDTA pH 8.0 containing 10% lysozyme and incubated at 37°C for 2 hours. Extraction of gDNA was performed using Wizard genomic DNA Kit (Promega). After quantification by nanodrop, the gDNA samples were sent to Maryland Genomics core at the University of Maryland School of Medicine. Following quality assessment with the Agilent 5200 Fragment Analyzer System, gDNA samples were subjected to pooled barcoded large insert (≥20 kb) library construction with size selection for HiFi Multiplex sequencing on the PacBio Sequel II platform. Complete, gap-free genomes of the four strains were generated using the PacBio SMRT 11.0.0 Improved Phased Assembler (IPA). Genome sequences were submitted to NCBI for PGAP annotation and released with accession numbers listed in Table S1. Table S1 also includes accession numbers of previous sequencing data from these strains.

## Acknowledgments

This work was supported by the Division of Intramural Research (DIR) of the National Institute of Allergy and Infectious Diseases and the Clinical Center, NIH.

